# Rapid growth and fusion of protocells in surface-adhered membrane networks

**DOI:** 10.1101/2020.03.10.980417

**Authors:** Elif S. Köksal, Susanne Liese, Lin Xue, Ruslan Ryskulov, Lauri Viitala, Andreas Carlson, Irep Gözen

## Abstract

Elevated temperatures might have promoted the nucleation, growth and replication of protocells on the early Earth. Recent reports have shown evidence that moderately high temperatures not only permit protocell assembly at the origin of life, but could have actively supported it. Here we show the fast nucleation and growth of vesicular compartments from autonomously formed lipid networks on solid surfaces, induced by a moderate increase in temperature. Branches of the networks, initially consisting of self-assembled interconnected nanotubes, rapidly swell into microcompartments which can spontaneously encapsulate RNA fragments. The increase in temperature further causes fusion of adjacent network-connected compartments, resulting in the redistribution of the RNA. The experimental observations and the mathematical model indicate that the presence of nanotubular interconnections between protocells facilitates the fusion process.

---

The important role of solid surface support for the autonomous formation of primitive protocells has been suggested earlier in the context of the origin of life^1,2^. Hanczyc, Szostak *et al*. showed that vesicle formation from fatty acids was significantly enhanced in the presence of solid particle surfaces consisting of natural minerals or synthetic materials^1,2^. Particularly the silicate-based minerals accelerated the vesicle generation.

In a recent report, we showed the autonomous formation and growth of surface adhered protocell populations as a result of a sequence of topological transformations on a solid substrate^3^. Briefly, upon contact with a mineral-like solid substrate, a lipid reservoir spreads as a double bilayer membrane. The distal membrane (upper -with respect to the surface-) ruptures and forms a carpet of lipid nanotubes. Over the course of a few hours, fragments of these nanotubes swell into giant, strictly unilamellar vesicular compartments. This relatively slow process is entirely self-driven and only requires a lipid reservoir as source, a solid surface, and surrounding aqueous media. The resulting structure consists of thousands of lipid compartments, which are physically connected to each other via a network of nanotubes. This formation process and the ability of these compartments to encapsulate ambient molecules, and to separate and migrate to remote locations, lead to the formulation of a new protocell hypothesis^4^. This addresses open questions about how primitive protocells might have formed and replicated on the early Earth, what exact physico-chemical mechanisms governed the growth and division of the membranes, and how cargo, *e*.*g*. RNA or other contents, was encapsulated and distributed. Prevailing hypotheses involving the self-assembly of amphiphiles in bulk aqueous medium explain the formation of protocells, but not the necessary subsequent steps, *e*.*g*. growth, replication, division, in a satisfactory manner.

In the former study, which was conducted at constant room temperature, the protocell nucleation and growth is a slow process occurring over the course of hours to days^3^. Under natural conditions, fluctuations in temperature are expected, the impact of which on the reported system has not been considered. Since the growth process is slow, the compartments often do not reach sizes large enough to establish physical contact in a reasonable time frame, and remain too far apart for fusion. Fusion has been considered a feasible means of protocell growth, a step required for self-proliferation^5^. In addition, the mechanical or osmotic stress on bilayer compartments can over time lead to the collapse of vesicular structures^6^.

In our current study we show that a temperature increase significantly accelerates the formation of membrane compartments and further initiates their fusion, which supports the recent findings of Jordan *et al*.^7^. The accelerated nucleation and growth lead to maturation of compartments, which eventually establish physical contact. The adjacent vesicular membranes fuse, resulting in redistribution of cargo, *e*.*g*. oligoribonucleotides. Nucleation and transformations strictly occur on the lipid nanotube networks and creates consistently and exclusively unilamellar membranous compartments. In addition to the experiments, we provide a finite element model which emphasizes that the presence of nanotubular connections between protocells facilitates the fusion. The findings can explain how protocells on the early Earth might have undergone rapid growth and replication, and provide new insight into our recently developed nanotube-protocell network hypothesis^4^.

## Results

### Enhanced protocell formation and growth

We deposited a multilamellar reservoir on a SiO_2_ surface. The reservoir spontaneously spreads on the surface in form of a circular double bilayer membrane^8^. The distal of the two stacked bilayers (upper bilayer with respect to the surface), ruptures due to continuous tensile stress^8^, resulting in formation of a network of nanotubes on the proximal bilayer^3^. Fragments of the nanotubes swell over time, and form unilamellar vesicular compartments. The autonomous transformation of lipid reservoirs into networks of surface-adhered protocells interconnected by lipid nanotubes, has been described in Köksal *et al*.^3^.

This precursor structure is a lipid nanotube network residing on a bilayer patch (**Fig. 1a**). Each liquid-filled nanotube has a cylindrical cross-section and consists of a single bilayer^3^ (inset to **Fig. 1a**). Next, we engage an IR-B (λ=1470 nm) laser to achieve a mild temperature increase in the vicinity of the membrane^9^(**Fig. 1b**). The IR radiation is applied through an optical fiber positioned by means of a mechanical micromanipulator on an inverted microscope (**S1, Fig. S1**). The position of the IR-laser fiber tip with respect to the lipid nanotube-covered membrane region is indicated by the yellow dashed lines in **Fig. 1c**. Due to the flat fiber tip, the laser radiation is not focused, but affects a cone-shaped water volume that extends to the solid surface. Details of the experimental setup are provided in the supplementary information (**Fig. S1**). The formation of giant unilamellar vesicles from disordered membrane layers induced by localized heating (>25 °C) has been reported before^10,11^. In our experiments, the thermal gradient leads to the instant formation and growth of vesicular compartments exclusively from the lipid nanotubes (**Fig. 1b**). The formed compartments are strictly unilamellar^3^ which is in contrast to the vesicles reported by Billerit *et al*.^10,11^ where a lamellarity distribution typical for swelling of stacked bilayers was observed. The majority of techniques for artificial vesicle generation feature this distribution^12,13^. **Fig. 1c-e** (**Movie S1**) show the laser scanning confocal microscopy time series of the process schematically described in **Fig. 1a-b**. Over the course of a few minutes of IR exposure, the unilamellar compartments form from the lipid nanotubes and rapidly grow (**Fig. 1d-e**). The fiber is positioned above the sample at a tilted angle, therefore the beam projects onto an ellipse-shaped area, clearly visible from the distribution of the protocells in **Fig. 1d-e**. The image series in **Fig. 1f-k** (**Movie S1**) demonstrate that the formed compartments are co-located with the nanotubes. During growth the compartments maintain their positions. In **Fig. 1l-m** the transformation from tube to spherical compartment, is schematically shown. Due to the temperature increase, the membrane viscosity is reduced and tension increases, leading to rapid inflow of lipids from low tension areas. Combined with a reduction of the high membrane curvature, this results in minimization of the surface free energy of the system^3^, which is the driving force for the transformation.

**Figure 1.**
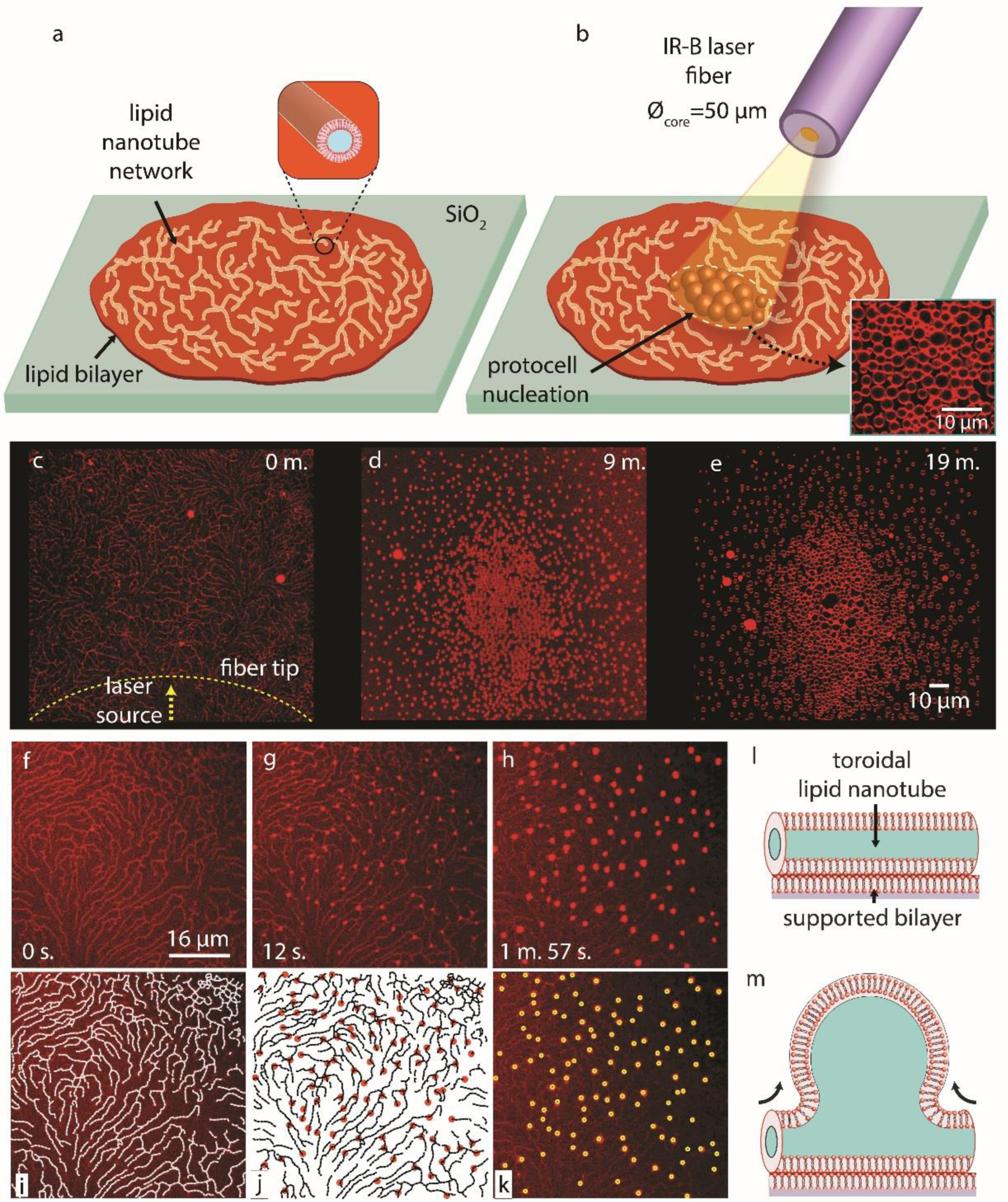
Heat-induced protocell formation from surface-adhered lipid nanotubes. **(a-b)** Schematic drawing summarizing the experiment. **(a)** network of hollow lipid nanotubes (inset) is residing on a SiO_2_-adhered bilayer. **(b)** rapid formation of protocells from the nanotubes as a result of mild heating. Inset on the lower right corner shows the confocal micrograph of protocells formed as a result of this process. All experiments have been performed in biological buffers. **(c-e)** laser scanning confocal microscopy time series of the process schematically described in **(a-b). (f-k)** confocal micrographs showing that the formed compartments and the nanotubes have co-localized. During growth, the compartments maintain their positions. **(i)** The outline of the nanotubes in panel **(f). (j)** The positions of the nucleation sites in **(g)**, indicated with the red circles, are superimposed on the network outlined in **(i). (k)** the positions of the nucleation sites in **(g/j)** are superimposed on the micrograph in **(h). (l-m)** Schematic drawing depicting the transformation from a nanotube to a vesicular compartment.

**Fig. 2a** (**Movie S1**) is a snapshot from a confocal time series of a heated membrane region, showing protocell growth. We created 31 elliptical rings (a quarter of each shown as a yellow dashed line) on the membrane and calculated the protocell densities, *i*.*e*., the number of protocells per area between two consecutive rings (Δ*Ar*_x_). An exception is the smallest ellipse at the centre which is considered as a whole. **Fig. 2b** is a plot of the protocell density in each ring versus the minor ellipse radius (*r*_x_) in **Fig. 2a**. The graph indicates that the protocell density increases with the temperature, which, due to the acceptance-cone of the fiber is gradually decreasing with distance from the center. The image series in **Fig. 2c-f** shows a membrane section decorated with lipid nanotubes, progressing to vesicular compartments. The confocal scans were recorded close to the proximal membrane (panel **e**), and across the equator of the vesicular compartments (panel **d**). Panel (**f**) is a 3D re-construction of the nanotube-adhered protocells in panels (**e-d**). For comparison, five selected compartments in the different panels of **Fig. 2e-f** are marked with numbers.

**Figure 2.**
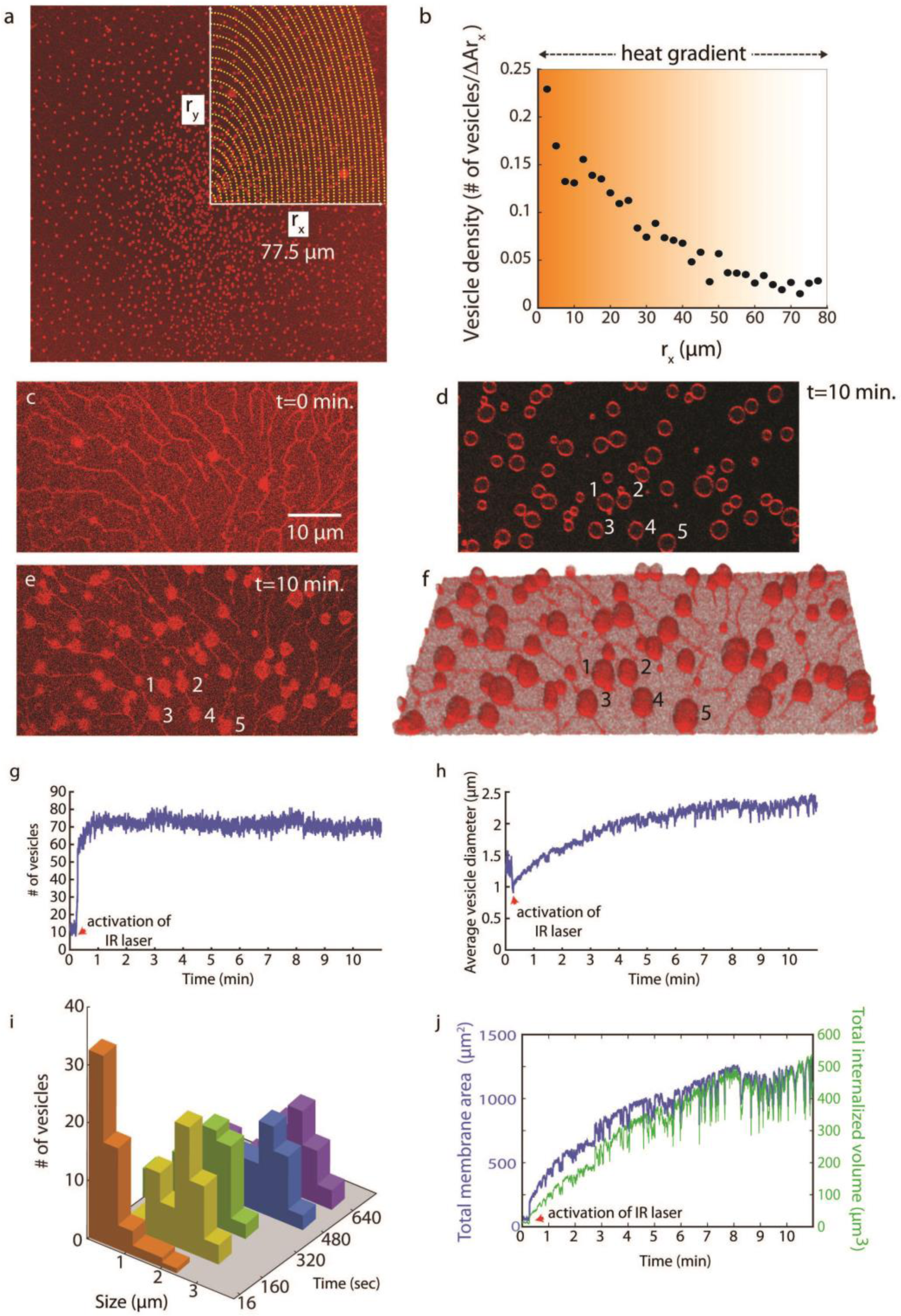
Characterization of protocell formation and growth induced by a mild heat gradient. **(a)** laser scanning confocal micrograph of a large membrane region with nucleating protocells. The area exposed to the IR laser is split to 31 hypothetical elliptical rings, the minor radius of which is expressed as *r*_x_ and the major radius, *r*_y_. A quarter of the outline of each ring is shown in yellow dashed lines. *r*_x_ of the outmost ring is 77.5 µm. **(b)** Plot showing the protocell density over distance *r*_x_. The protocell density is calculated as the number of protocells in each individual elliptical ring. **(c-f)** Confocal micrographs of a nanotube network leading to nucleation and growth of protocells exposed to heat gradients for 10 min.0020(**c**) nanotube network before local heat exposure **(d)** cross section of protocell sample from the equator after heat exposure (top view) **(e)** cross section of sample close to the surface after heat exposure (top view) **(f)** 3D reconstruction of the formed protocells. Plot showing **(g)** the number and **(h)** the average diameter of the protocells formed in **(c-f)** over 10 min. **(i)** The histograms depicting the size distribution of protocells over time. Each color represents the size distribution at a given time point. **(j)** Plots showing total membrane area and total membrane volume of the protocells in **(c-f)** during their formation and growth.

We used a micro-thermocouple to determine the temperature of the affected region in the experiments. The details of the measurement have been provided in the Supplementary Information **(S2, Fig. S2)**. The estimated temperatures are 40-90 °C. ∼40 °C leads to rapid nucleation, and ∼70 °C to rapid growth. ∼90 °C results in fusion of the compartments, which will be described in detail below.

For the membrane section depicted in **Fig. 2c-f** (**Movie S3**), the total count of the protocells versus time is shown in **Fig. 2g**, and the average protocell diameter versus time in **Fig. 2h**. The compartments nucleate instantly with activation of the IR-laser. Their number remains constant (**Fig. 2g**), while their diameter is increasing (**Fig. 2h**). The size distribution of the protocells at five different time points throughout the experiment is shown in **Fig. 2i**. Small protocells (*d* < 1 µm) dominating at the early stages (orange histogram), later evolve into larger protocells (*d* > 3 µm, blue and purple histograms). The development of the membrane area and the internal volume of the protocells during the experiment (**Fig. 2c-f**) is shown in **Fig. 2j**. The total membrane area was approximately 1250 µm^2^ at the end of the experiment. This corresponds to a nanotube of 4 mm length (*Ø* = 100 nm). We do not observe such a high tubular density in **Fig. 2c**. The membrane material forming the protocells therefore likely originates from a remote membrane reservoir and migrates through the nanotubes^14,15^. A comparison between panel (**c**) and (**e**) reveals that the majority of the nanotubes remain intact. This means that they are not the major source of protocells formed in the process. Provision of the membrane material through the proximal bilayer is in principle also possible, but we have earlier presented the argument that this is rather unlikely^3^.

### Protocell fusion

We observe that the temperature increase further induces fusion of adjacent compartments. **Fig. 3a-b** shows a membrane region in which rapid merging was observed. In **Fig. 3a** the protocells are shown before fusion, and in **Fig. 3b** after fusion. The areas in which fusion of two or more compartments occurs, are encircled in white dashed lines and numbered (panel **a**). The merged compartments are represented in **Fig. 3b** by the same numbers. **Fig. 3c** shows the number (orange graph) and average diameter (blue graph) of the protocells shown in **Fig. 3a-b** over time (*cf*. **S3** for details of the image analyses). The total number of compartments decreases, and the average diameter increases accordingly (panel **c**). The micrographs in **Fig. 3** (**Movie S4** and **S5**) show different surface regions of the same confocal microscopy recording. The black arrows in panel **c**, labeled with panel names, indicate the time points at which the corresponding images were recorded.

**Figure 3.**
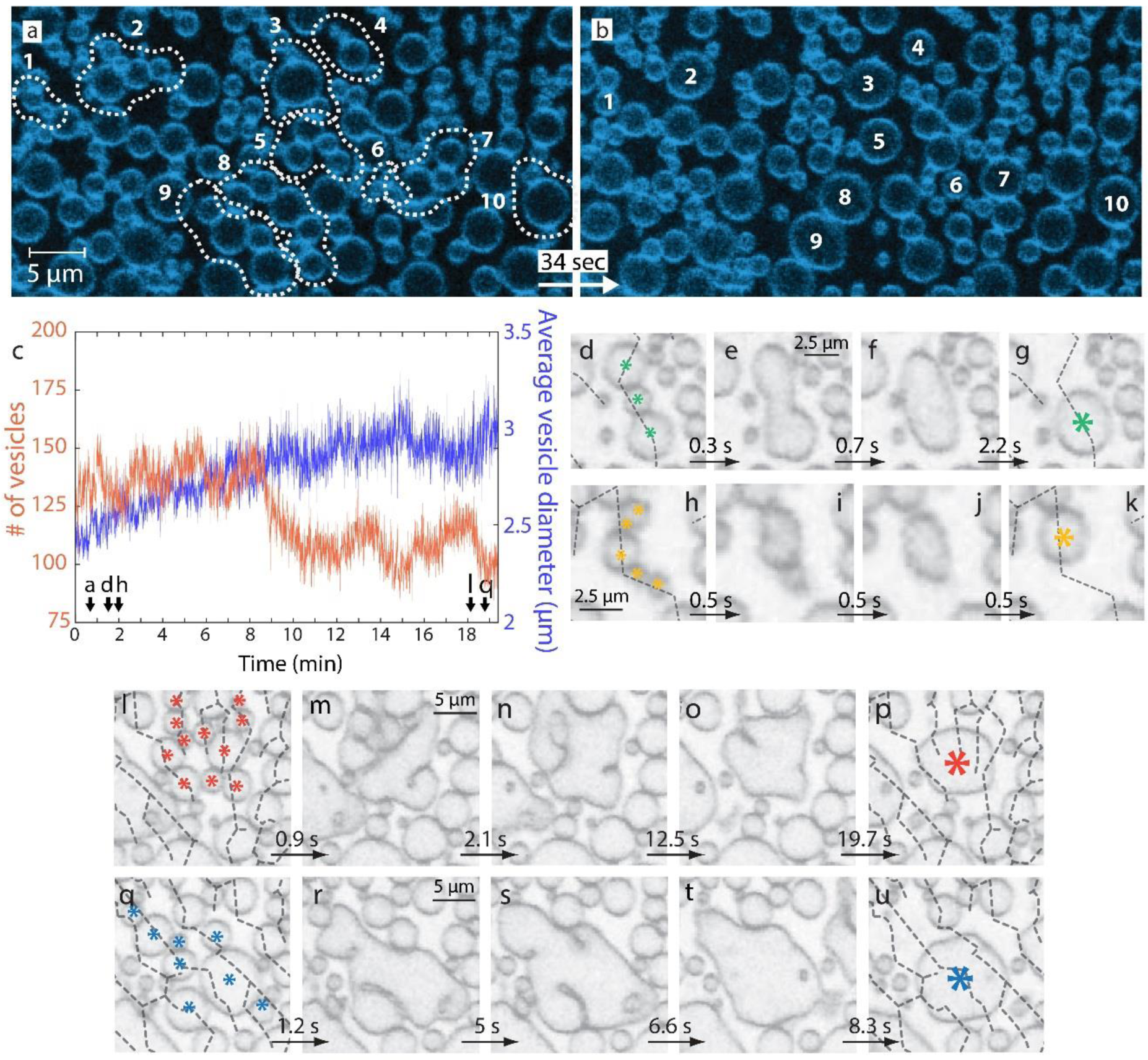
Heat induced protocell fusion. **(a-b)** Confocal micrographs showing the fusion of several protocells formed out of a lipid nanotube network upon exposure to the IR laser. (**a**) The group of protocells which later merge after exposure to heat gradient, are encircled with dashed lines. Each encircled region is numbered. (**b**) The merged protocells. Each protocell has been formed or grown as a result of the fusion of the multiple, originally separated protocells shown in (**a**) The group of protocells and their fused version are numbered identically in (**a**) and (**b**). **(c)** Plots showing the number of protocells (orange graph) and average protocell diameter (blue graph) over the complete course of the experiment partly shown in (**a-b**). **(d-k)** Protocell fusion on same nanotube. (**d-g**) and (**h-k**) show two different fusion events in which the protocells on the same nanotube rapidly merge. **(l-u)** Fusion of protocells which are originally located on separate nanotubes. (**l-p**) and (**q-u**) show two different events during which vesicular compartments, originally located on different nanotubes, later fuse.

We explored whether fusion events only occur among the compartments residing on the same nanotube, or if the fusion between compartments on different nanotubes is possible. We therefore investigated membrane regions of known network topology. **Fig. 3d-g** and **3h-k** shows the fusion of two different sets of compartments, each set residing on the same nanotube (black dashed lines). Three compartments marked with green asterisks in **Fig. 3d** merge into a single protocell within a few seconds (panels (**e-g**)). In **Fig. 3h-k**, five protocells (yellow asterisks) merge. In both cases, the fusing protocells reside on the same nanotube. If protocells located on different nanotubes grow and eventually establish physical contact, we also observe fusion events. **Fig. 3l-p** (red asterisks) and **3q-u** (blue asterisks) depict two examples. The original positions of the nanotubes in each recording are indicated by black dashed lines (**Fig. 3l** and **3q**).

### Mechanism of fusion

Increase in temperature results in an increase in fluidity of the membrane, leading to the rapid fusion of the initially distinct, adjacent membranes^16,17^.Upon fusion, the membranes relax to a form that minimizes the membrane energy. The transformation from two small containers to a single large one reduces the curvature, while the membrane area is maintained. In our experiments, fusion of nanotube-connected lipid compartments can occur in two different ways: it either begins near their equator, *i*.*e*., where their lateral extension is the largest and the compartments touch first (**Fig. 4a**), or they fuse at the base, mediated by the connecting membrane nanotube (**Fig. 4b**). In order to determine which of these two scenarios is energetically the most favorable, we performed a set of numerical finite element simulations^18,19^ (*cf*. **S4** for details). Since the thickness of the lipid bilayer (∼ 5 nm) is much smaller than the typical size of the membrane tube (> 100 nm) and the attached compartment (> 1 µm), we treated the membrane as a thin elastic surface. For the simulation, we considered two adjacent vesicular compartments of the same size, where the compartments share a surface-adhered membrane tube. The edges of the numerical domains were defined by open nanotubes which were restricted to form a cylinder of radius *r*_t_. The tube length was set to 15 *r*_t_ and the total membrane area to 450 *r*_t_^*2*^. In dimensional units this corresponds to two spherical compartments, each with a diameter of ∼0.4 μm, connected through a tube with a diameter of ∼100 nm.

**Figure 4.**
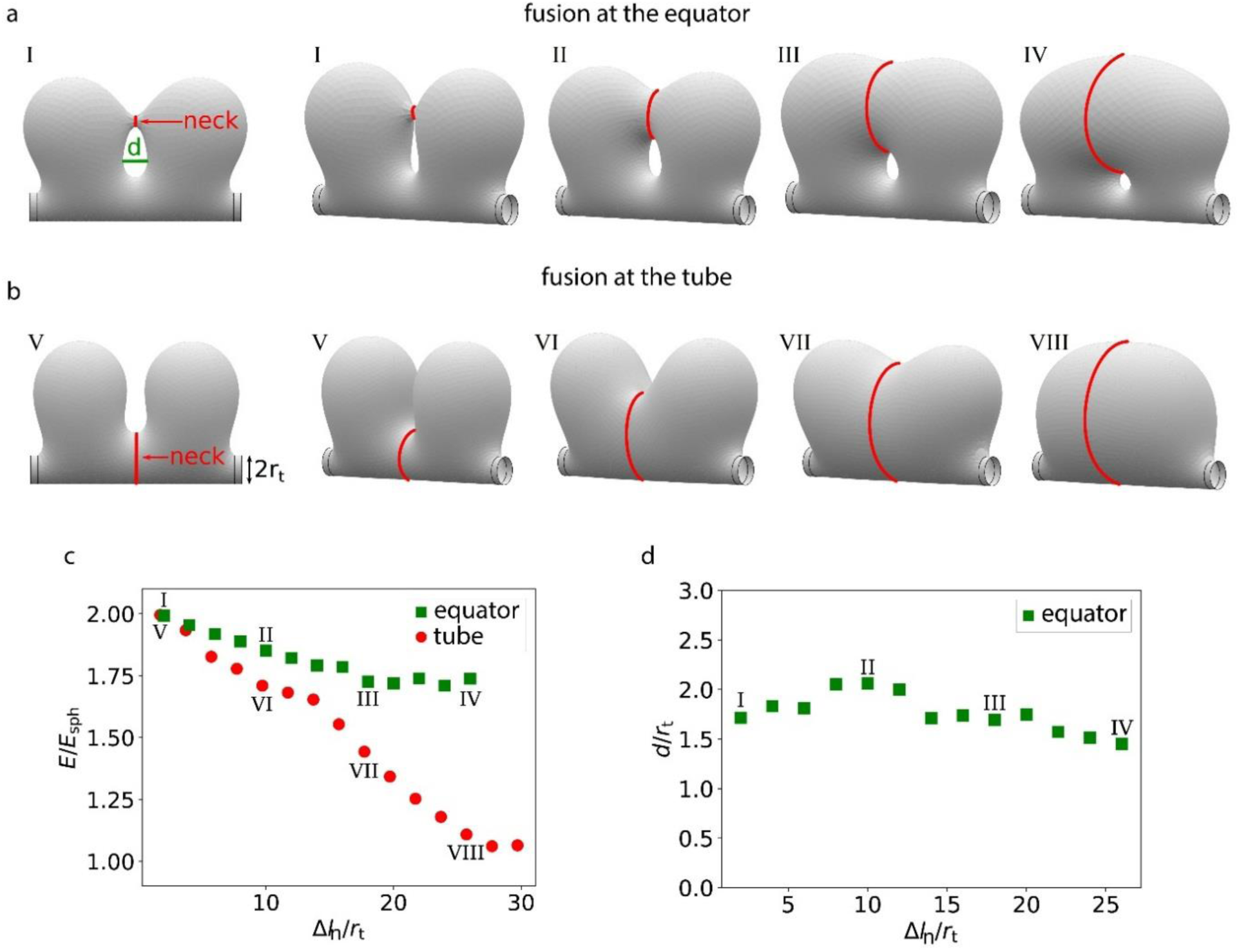
Mathematical model for fusion. Two compartments of equal size are connected to a membrane tube with diameter 2*r*_t_. Simulation snapshots are shown as the compartments fuse either at the compartments’ equator region (**a**) or at the connecting tube (**b**). The outer left snapshot on (**a**) and (**b**) show a side view parallel to the membrane tube, while the other snapshots are tilted to better illustrate the expansion of the fusion neck. (**c**) The bending energy *E*, rescaled by the bending energy of the spherical compartment *E*_sph_, decreases as the length of the contact line *Δl*_n_ increases. (**d**) If the compartments fuse initially at their equator, a cavity forms between the fusion site and the membrane tube, with a stable diameter *d* that is similar to the diameter of the membrane tube.

We keep the membrane area in the simulations constant such that the membrane shape and energy are solely determined by the minimization of the bending energy. In the initial configuration, the two compartments either form a fusion pore (neck) near their equator (**Fig. 4a**), or they fuse by consuming the nanotube (**Fig. 4b**). In both cases the bending energy of two well separated compartments, *i*.*e*. narrow neck with small *Δl*_n_, is similar to the energy of two independent spherical compartments. In the latter case, we consider the portion of the tube that connects the two compartments, the neck region. In each simulation the circumference of the neck is kept constant, while the position of the neck is free and hence determined by the energy minimization. **Fig. 4c** shows the development of the bending energies for fusion initiated at the equator (squares) and at the tube (circles), as we systematically increase the neck circumference *Δl*_n_. We obtain the bending energy, according to the Helfrich theory^20^ by integrating the square of the mean curvature *H* over the membrane surface area *A*: 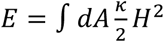, where *κ* is the bending rigidity. The energy is normalized by the bending energy of a spherical vesicle, *E*_sph_=8π*κ*. If the vesicle fusion starts at the tube, we observe an increase in neck circumference *Δl*_n_. Note that in this case the initial circumference 2π*r*_t_ is subtracted. As the neck expands, the vesicles fuse and the surface free energy decreases. For vesicle fusion at the base, the energy approaches that of a single spherical vesicle. In contrast, for vesicles fusing at the equator, the energy reaches a plateau that is about 75% larger than the energy of a spherical vesicle. If the vesicles start to fuse at their equator, a circular pore forms between the fusion site and the membrane tube. Our simulations show that the pore stabilizes with a diameter (IV in **Fig. 4a**) similar to the diameter of the membrane tube (**Fig. 4d**). The fusion process should predominantly start at the tube, since this scenario is energetically more favorable and allows for complete fusion of the two vesicles (*cf*. **S4** for model, **S5** for corresponding experimental observations).

### Encapsulation and re-distribution of RNA upon fusion

In order to investigate the merging of the contents of fusing compartments, we employed an open-space microfluidic pipette^21^ and loaded several compartments with fluorescently labeled RNA oligomers (**Fig. 5a**). Open-space microfluidic delivery is an effective means to create a local chemical environment. Subsequently applied mild heating led to the fusion of the RNA-loaded protocells with adjacent, initially unloaded compartments, and the RNA oligomers were redistributed (**Fig. 5b**). **Fig 5c** (**Movie S7**) shows the loading process: a population of surface-adhered protocells in the recirculation zone of the microfluidic device, dispensing a solution containing fluorescent RNA fragments (top view). The border of the recirculation zone is marked with white dashed lines (**Fig. 5c**). Initially, the protocells appear as black dots within the recirculation zone, since they only contain buffer (**Fig. 5c**). Over time, some of the protocells encapsulate the RNA fragments (**Fig. 5d**). Encapsulation of water-soluble fluorescein inside the surface-adhered protocells was shown earlier in a similar experiment at ambient temperature^3^, and explained by the involvement of transient pores of sufficient size and stability. We observe that the encapsulation efficiency of RNA oligomers is significantly smaller than of fluorescein (*cf*. **S6** for comparative experiments). We attribute this to both the size and the charge differences between the molecules. **Fig. 5e-m** is a sequence depicting how the RNA fragments are redistributed during fusion. **Fig. 5c-g** shows both the membrane and the RNA fluorescence emission channels, **Fig. 5h-j** the membrane, and **Fig. 5k-m** the RNA. **Fig. 5n-p** shows the fluorescence intensity along the lines indicated with white dashed arrows in **Fig. 5k-m**, respectively. The initially separate signals in **Fig. 5n** merge into a signal of lower intensity (**Fig. 5o-p**).The decrease may be due to two reasons. Upon activation of the IR laser the fluorescence intensity is reduced^22^. It is also possible that some of the internal contents is lost by leakage. The closure of the pores and retaining of the bulk of the encapsulated material is consistent with our earlier experiments of ∼3 orders of magnitude longer duration (seconds vs. hours). **Fig. 5r-t** (**Movie S7**) shows snapshots of the fusion process of three compartments. In **Fig. 5s** depicts the diffusion of fluorescent fragments during shape optimization of merging containers.

**Figure 5.**
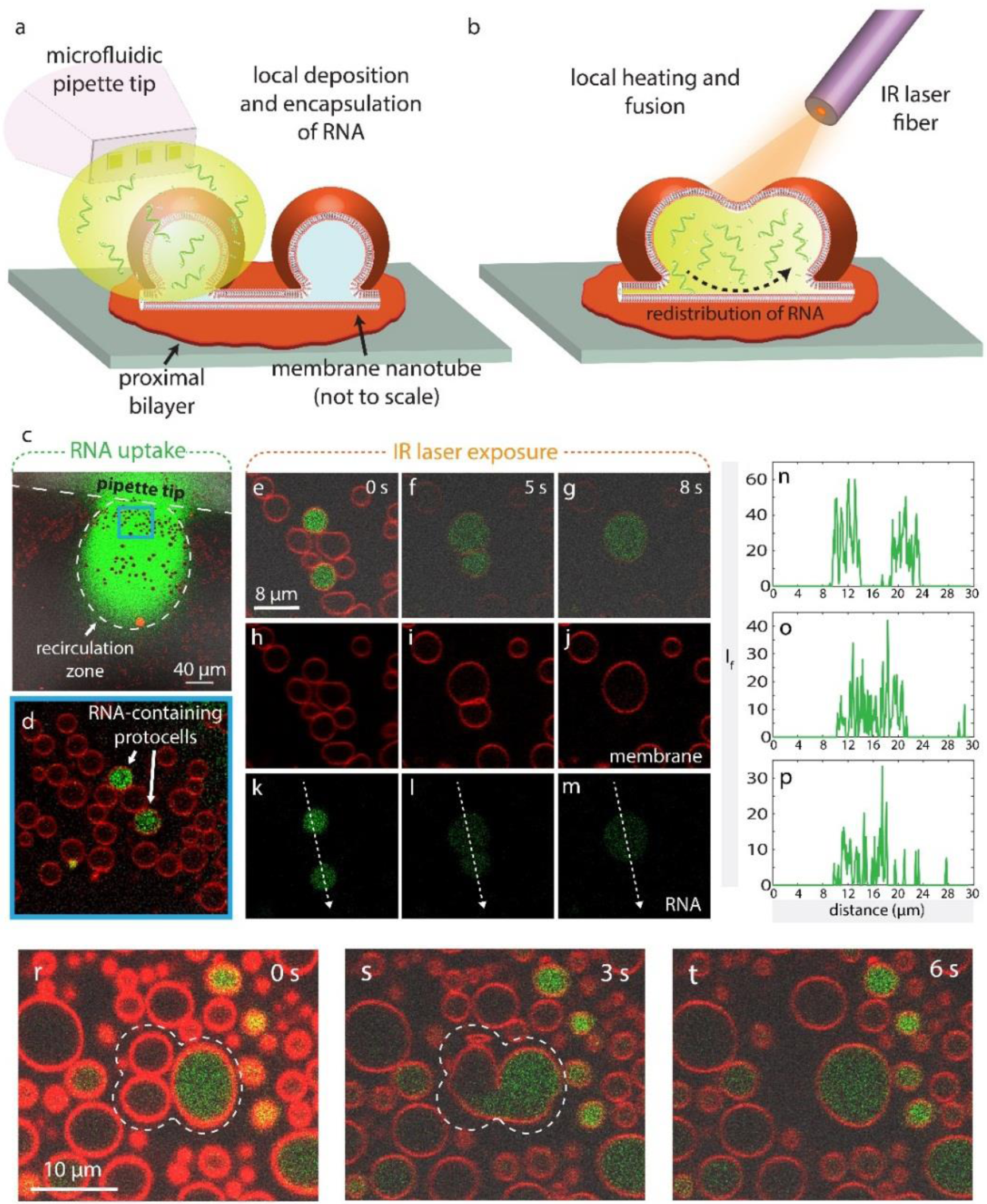
RNA encapsulation and redistribution. **(a-b)** Schematic drawing showing the experimental setup. **(a)** An open-space microfluidic device is used for the superfusion of RNA-oligonucleotides with a designated membrane area populated with protocells. **(b)** IR laser is activated to induce fusion, leading to the redistribution of pre-encapsulated RNA into the fused protocell. **(c-d)** RNA uptake. **(c)** Confocal micrograph of a membrane area with the microfluidic pipette re-circulating RNA above it (top view). The protocells in the recirculation zone appear as black dots. **(d)** Magnified view of the blue frame in **(c)** after termination of recirculation. Two protocells contain RNA. **(e-m)** Laser scanning confocal microscopy images showing the fusion of RNA encapsulating protocells and redistribution of contents upon fusion. **(e-g)** Membrane, RNA fluorescence and bright field channels are overlaid. **(h-j)** membrane fluorescence channel only. **(k-m)** RNA fluorescence channel only. **(n-p)** Plots showing the fluorescence intensity over the white dashed arrows in **(k)**,**(l)** and **(m)**, respectively. **(r-t)** Laser scanning confocal microscopy images showing the fusion of RNA-encapsulating protocells and redistribution of contents upon fusion. Membrane, RNA fluorescence and bright field channels are overlaid.

## Discussion

### Protocell nucleation sites

**Fig. 2g** reveals that, once the protocells nucleate, their total number remains constant during growth. This indicates that the sites of the nucleation are pre-determined and nucleation is enhanced by the increased temperature. In **Fig. 1j** the locations of the nucleation appear to coincide with Y- and V-junctions^23^ on the nanotubes. Such membrane topologies are caused by pinning, *i*.*e*., simultaneous binding of Ca^2+^ to multiple lipid headgroups, which is facilitating the cohesion between two stacked bilayers ^8,24,25^ or between a bilayer and a solid interface^8,26,27^. In a previous study, the transformation of a Y-junction to a small vesicle, due to chemical chelator-induced de-pinning of Ca^2+^, has been already shown^23^. In the current study, the Ca^2+^ de-pinning and reversal of membrane adhesion is not caused by chelators, but is due to the temperature increase^9^. The compartments are also observed exclusively at junction points (**Fig. 1j**).

### Mechanism of rapid growth and fusion

The main driving force for the transformation of the nanotubes to protocellular compartments is the minimization of membrane curvature. The natural growth process in the previously reported system was slow (∼h)^3^. The membrane replacement rate for the spontaneous inflation of a tube to a 5 μm vesicle, was estimated the to be ca. 2×10^−3^ µm^2^/s^3^. In contrast, in the current study the transformation occurs within minutes. We estimate the replacement rate to be 2 µm^2^/s, about three orders of magnitude higher than observed at room temperature. We attribute the facilitated protocell growth to the enhanced ability of lipid material to flow to the area of nucleation, due to the temperature increase in that area. The locally elevated temperature causes an increase in the membrane fluidity and in the membrane tension in the affected area. The tension increase causes Marangoni flow of lipids in the surrounding membrane region with relatively low membrane tension, towards the heated membrane region with high tension. The result is the rapid growth of previously nucleated vesicular buds to cell-sized unilamellar compartments, some of which eventually establish physical contact with each other.

Fusion of lipid compartments that are in close proximity does not occur spontaneously, but requires external stimuli. There have been several studies focusing on the fusion of giant amphiphile vesicles as model systems of proto- or contemporary cells. The reported fusion mechanisms vary. Some examples are the fusion driven by attraction of oppositely charged vesicles^28^, fusion induced by multivalent ions between vesicles of special amphiphilic compositions, *e*.*g*. Eu^3+29^ or La^3+30^, ultraviolet light radiation-induced fusion ^31^, electrofusion^32^, fusion involving amphiphilic catalysts^33^, also in combination with thermal cycles and pH changes^34^, and the fusion mediated by the hybridization of complementary SNARE proteins^35^ or DNA linkers ^36^, embedded in initially distinct vesicle membranes.

The vesicular membranes we utilize in this study do not contain embedded species which would facilitate fusion. The membranes and their individual monolayer leaflets possess the same composition, thus they are free of spontaneous curvature and have the same electrostatic potential. All experiments are performed at constant pH, under identical conditions. The main stimulus for fusion is the controlled increase in temperature. The temperature increase is known to lower the microviscosity of the membrane and facilitate the formation of defects, especially in the presence of multivalent ions^37,38^. Ca^2+^ binding perturbs the membrane by pulling the headgroups inwards^24,37^, causing formation of defects. With localized heating this process is facilitated^38^, resulting in fusion^39^.

### Impact of nanotubes in growth and fusion

**Fig. 3** and **Fig. 4** shows that the nanotubes, which physically connect distant protocells, facilitate fusion. During growth, the nanotubes can provide an additional advantage for compound delivery. They provide a transport pathway for a continuous influx of molecules through the network, by which larger molecules can potentially also be transported. The transport of molecules or particles through nanotubes occurs by diffusion ^40,41^, or is tension-driven (Marangoni flow)^14,15^. The transport phenomena within the nanotube networks have not been investigated here, but the earlier established evidence of nanotube-enhanced transport between lipid vesicles combined with the involvement of nanotubes in the fusion process, as elucidated in this study, points to a beneficial contribution of an existing tubular network for growth, transport and fusion of protocells.

### Impact of temperature in the context of origin of life

In this study we show that a successive increase in temperature from 20 °C to approximately 40, 70 and 90 °C on a nanotube network facilitates the nucleation, growth and fusion of surface adhered protocells. The role of temperature has been a central discussion point in the origin of life debate ^42^. The competing hypotheses regarding the environment for the emergence of the RNA world, concentrate either around deep ocean hydrothermal vents, or around warm ponds^42^. A major criticism for the emergence of life in hydrothermal environment^43^ has been the hot temperatures, large pH gradients, high salinity and high concentrations of divalent cations, which may adversely affect the amphiphile compartment formation. The hot environments typically referred to in such discussion involve black smoker type hydrothermal vents where temperature can commonly exceed 300 °C. In 2000, a new type of hydrothermal vent: the *Lost City hydrothermal field* (LCHF) with a chemical composition similar to lavas that erupted into the primordial oceans on early Earth, was discovered^44^. The temperature range of the LCHF is 40-90 °C, surprisingly similar to the experimental conditions used in this work that promote compartment formation, growth and fusion. This temperature range also represents the conditions in warm ponds: 50-80 °C^42^. Recent evidence shows that the mixtures of single chain amphiphiles form vesicles most readily at temperatures of ∼70 °C in aqueous solutions containing mono- and divalent cations in broad pH range^7^. In the light of these observations, it appears that warm temperatures of ponds or LCHFs can allow and even favor protocell compartmentalization. Our investigation focusing on the subsequent steps, *i*.*e*. the rapid growth and fusion, is in alignment with these recent findings.

Apart from temperature, another point disfavoring the hydrothermal vent hypothesis over the warm pond hypothesis, has been the lack of dry-wet cycles, which is known to significantly facilitate polymerization, *e*.*g*. from nucleotides to RNA^42^. In our experiments, the lipid reservoirs, *i*.*e*. multilamellar vesicles (MLVs), from which the double bilayer films spread and proceed to protocell formation, are the product of a dry-wet cycle. The lipid layers form in a dry environment and upon hydration they spontaneously form MLVs. It is conceivable that this is a repeatable process. Accordingly, protocell formation, growth and fusion events we report here can in principle occur during dry-wet cycles.

## Conclusion

We show that the nucleation, growth and fusion of protocells are significantly accelerated and enhanced at temperatures ranging from 40 to 90 °C. Some of the protocells generated in this manner have been demonstrated to encapsulate RNA, and to redistribute it upon fusion with other compartments. In the context of protocell development on the early Earth, these results suggest that both *Lost City*-type hydrothermal vents, and warm ponds could have been a suitable environment for protocell formation, growth and fusion events. Additionally, a supporting surface in conjunction with the physical interconnections provided by the spontaneously formed nanotubular networks pose an advantage over lipid assemblies in bulk solution. Neighboring vesicles can join and fuse more rapidly than in bulk suspensions, where protocells would only randomly encounter each other for limited periods of time. To what extent it is possible for emerging protocells to chemically communicate prior to, and during, fusion processes through interconnecting tubes remains to be elucidated. If this can be verified, new hypotheses for primordial chemical transformations within primitive membrane structures in the early Earth environment can be experimentally investigated.

## Materials and Methods

### Surface fabrication & characterization

A ∼84 nm SiO_2_ film was deposited onto Menzel Gläser (rectangular) or Wilco Well (circular) glass substrates by either E-beam, or thermal Physical Vapor Deposition, using an EvoVac (Ångstrom Engineering) or L560K (Leybold) evaporator. The thickness of the films was verified by ellipsometry (SD 2000 Philips). No pre-cleaning was performed before deposition. The substrates were stored at room temperature prior to use.

### Formation of lipid nanotube network and protocells

The lipid nanotube network on a solid supported bilayer was formed as described earlier^3^. Briefly, a stock suspension of multilamellar lipid reservoirs containing 50% soybean polar lipid extract, 49% *E. coli* polar lipid extract and 1% Rhodamine-PE or Cy5-PE were prepared by the dehydration/rehydration method^45^. An aliquot of 4 µl from this suspension was dehydrated in a desiccator for 20 min. The dry film was rehydrated with ∼1 ml of HEPES buffer containing 10 mM HEPES and 100 mM NaCl, pH 7.8, for 10 min to form multilamellar reservoirs. The reservoirs were then transferred into an open-top observation chamber on a SiO_2_ substrate. The chamber contained ∼1 ml of HEPES buffer with 10 mM HEPES, 100 mM NaCl and 4 mM CaCl_2_, pH= 7.8. On the SiO_2_ substrate the reservoirs self-spread as a double bilayer. The distal bilayer ruptures^8^, and a nanotubular network forms on the proximal bilayer^3^. Protocells on nanotubes were either formed spontaneously overnight (RNA redistribution experiments) or within seconds or minutes using local IR-B radiation (nucleation, growth and fusion).

### Heating system

The lipid-nanotube network was heated locally using IR-B laser radiation through a flat optical fiber tip. A 1470 nm semiconductor diode laser (Seminex) in combination with a 50 µm core diameter, 0.22 NA multimode optical fiber (Ocean Optics). The fiber was prepared by removing the outer sheath cladding, followed by carefully cutting and polishing using a fiber cleaning kit (Ocean Optics). The fiber was positioned using a 3-axis water hydraulic micromanipulator (Narishige, Japan) and the tip was located at 50 µm from the surface, resulting in a volume of approximately 1 nL being efficiently heated. Three different laser intensities were employed. The laser current was adjusted to 0.72 A (protocell nucleation), 0.97 A (growth) and 1.21 A (fusion). The temperature was determined directly by a micro-thermocouple *in-situ* (*cf*. supplementary information S2 for details).

### Encapsulation with microfluidic pipette

An open-volume microfluidic device/pipette ^21^ (Fluicell AB, Sweden), positioned using a second 3-axis water hydraulic micromanipulator (Narishige, Japan), was used to expose the matured surface-adhered protocells to Ca^2+^-HEPES buffer containing 100 µM fluorescein sodium salt (Sigma Aldrich), 40 µM FAM-conjugated RNA oligonucleotides (Dharmacon, USA), at pH 7.8.

### Microscopy imaging

A confocal laser scanning microscopy system (Leica SP8, Germany), with an HCX PL APO CS 40x (NA 1.3) oil objective was used for acquisition of the confocal images. The utilized excitation/emission wavelengths for the imaging of the fluorophores, were as follows: λ_ex_: 560 nm, λ_em_: 583 nm for membrane fluorophore Rhodamine-PE, λ_ex_: 655 nm, λ_em_: 670 nm for Cy5, λ_ex_: 488 nm, λ_em_: 515 nm for fluorescein (SI), λ_ex_: 494 nm, λ_em_: 525 nm for FAM.

### Image processing/analysis

3D fluorescence micrographs were reconstructed using the Leica Application Suite X Software (Leica Microsystems, Germany). Image enhancements to fluorescence micrographs were performed with the NIH Image-J Software and Adobe Photoshop CS4 (Adobe Systems, USA). Schematic drawings and image overlays were created with Adobe Illustrator CS4 (Adobe Systems, USA). Protocell counts, density, size distribution, total membrane area and volume analyses were also performed in Image-J and plotted in Matlab R2018a. The analysis of protocell number and size over time during fusion was performed with Matlab. Fluorescence intensity profiles were drawn in Matlab after applying median filtering.

## Supporting information

Koksal_etal_SI

Sample Matlab Script

## Acknowledgements

We thank A. Jesorka from Chalmers University of Technology, Sweden, for technical advice on temperature measurements. This work was made possible through financial support obtained from the Research Council of Norway (Forskningsrådet), Project Grant 274433, UiO: Life Sciences Convergence Environment, the Swedish Research Council (Vetenskapsrådet), Project Grant 2015-04561, as well as the startup funding provided by the Centre for Molecular Medicine Norway (RCN 187615), and the Faculty of Mathematics and Natural Sciences at the University of Oslo. S.L. and A.C. gratefully acknowledge funding from the Research Council of Norway Project Grant 263056.

## Author Contributions

E.K and I.G. designed the research, E.K., S.L., L.X., R. R, A.C. and I.G. performed the research, E.K., L.X., R. R, L.V. analyzed the experimental data, and all authors contributed to the drafting of the paper. I.G. suggested the investigation of the behavior of lipid compartments in thermal gradients and supervised the project.

