## Supplementary material for "Rapid growth and fusion of protocells in surface-adhered membrane networks": Koksal_etal_SI

**This PDF file includes:**

Supplementary text
- S1. Experimental setup
- S2. Characterization of IR-laser heating
- S3. Image analyses
- S4. Details of computational model
- S5. Stable pore formation in fused compartments
- S6. Encapsulation of fluorescein vs. RNA by the protocells

Figures S1 to S7
Captions for movies S1 to S7
References for SI

**Other supplementary materials for this manuscript include the following:**

Movies S1 to S7
Matlab Script: Detection of vesicular compartments from micrographs

### S1. Experimental setup

The optical fiber for co-application of IR-B radiation and the microfluidic pipette for the superfusion of RNA-oligonucleotides were positioned above the substrate surface, using 3-axis water hydraulic micromanipulators (**Fig. S1**). The tip of the fiber and the pipette are placed on opposite sides in order to target the same membrane area on the substrate.

The inset in the upper left corner of **Fig. S1** shows the flat polished tip of the optical fiber with a core diameter of 50 µm.

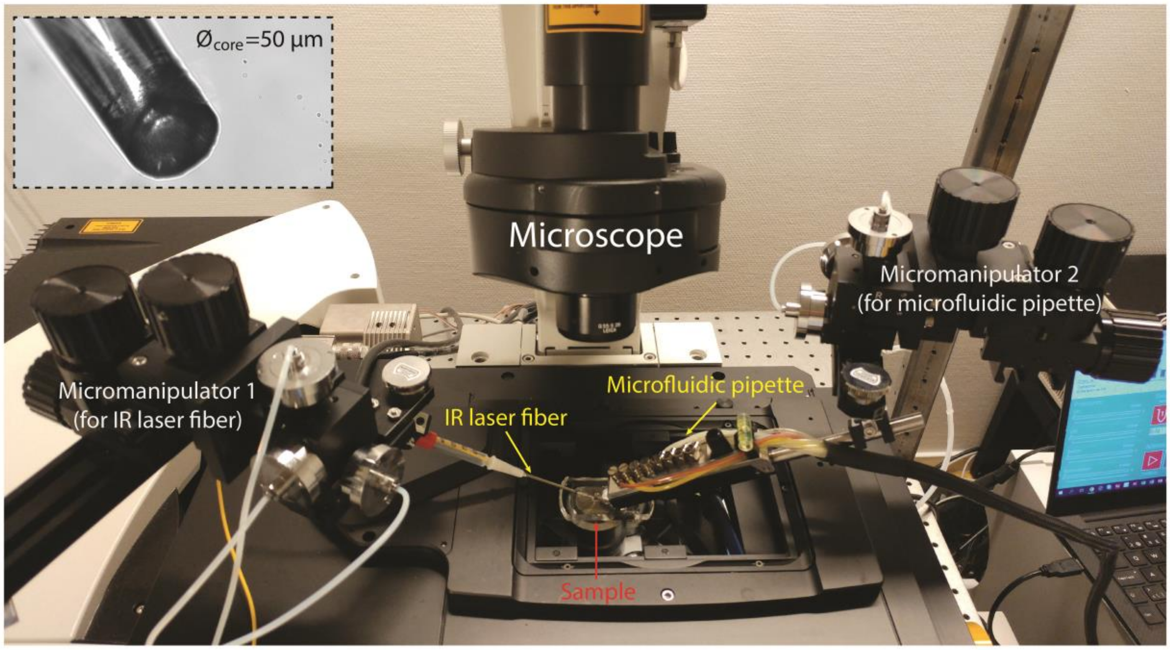


**Figure S1 | Photograph of the experimental setup**. Inset: flat tip of the IR laser fiber.

### S2. Characterization of IR-laser heating

The local temperature changes caused by the IR laser heating were estimated from the absolute temperature measurements with a CHCO-005 E-type microtemp thermocouple (junction diameter: 25 µm, Omega Engineering, UK). Optical fiber and thermocouple were positioned above the sample using motorized micromanipulators (Scientifica, UK) (**Fig.S2a**). For the measurements, the thermocouple was placed above the surface in the center of the heated region. Temperatures were recorded while the thermocouple was lifted up in z-direction in 10 µm steps. Measurements were performed for three different laser intensities, as employed in the compartment formation, growth and merging experiments. Plots of the obtained data are presented in **Fig. S2b**.
Due to the unavoidable direct absorption of IR irradiation by the thermocouple itself, the measured temperatures are somewhat higher than the actual temperatures in the medium. This is particularly grave in the range between 0-40 µm in z-direction (open circles in **Fig. S2b**), where the measured temperatures increase to unphysical values above the boiling point of water. Since boiling is not observed, we can conclude that the actual temperatures are below 100 °C. These data points were excluded, and only the data points shown with filled circles are fitted by means of a linear regression in order to estimate the temperature on the surface (*z*=0). According to this approximation, the lipid sample is heated up to temperatures of 39, 89 and 122 °C (as measured, see section

below for discussion of these values) when the laser diode current is 0.73, 0.97 and 1.22 A, respectively. The directly measurable laser current is stated rather than the laser power, which can be extracted from the U/I/P chart in the manufacturer data sheet (www.seminex.com) of the 4PN-104 1470 nm (NA 0.22 fiber coupled) laser diode. The continuous wave power output at 1 A corresponds to approximately 100 mW.

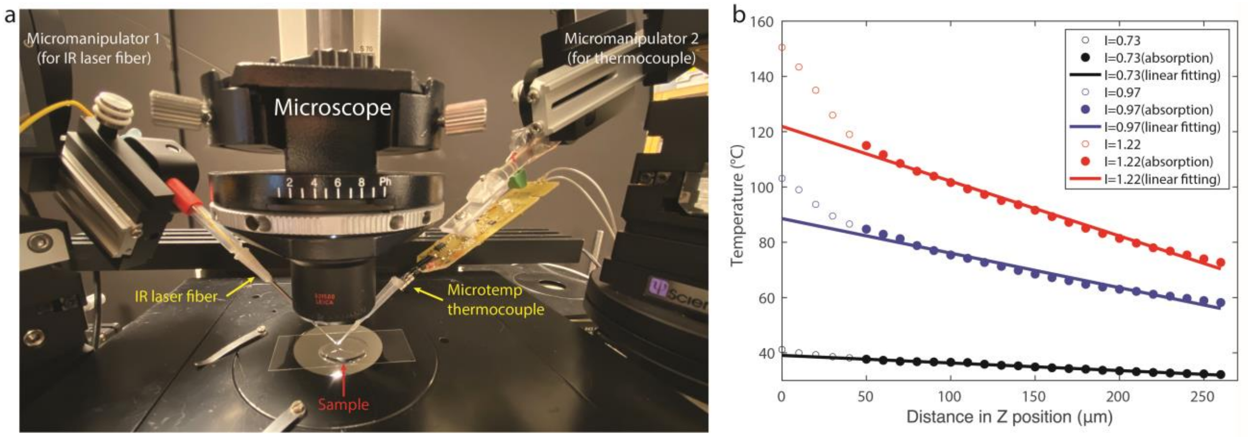


**Figure S2 | Characterization of the local temperature profile caused by IR-laser heating. (a)** Photograph of the experimental setup for the temperature measurement using thermocouple. **(b)** Plot of measured temperatures at three different laser intensities employed in the experiments.

We can distinguish between three different temperature ranges for each of the three different laser settings. There may be an additional component in the temperature/distance relationship for the two higher laser power settings for the height range between 200 and 250 µm distance from the surface, which corresponds to the situation where the thermocouple is outside the irradiated volume (cone of acceptance), which we have not considered in the analysis. At the lowest power setting, the direct absorption by the thermocouple metal is greatly reduced, due to the almost complete absorption of the light by the water volume in between fiber end and thermocouple. The measured values are for this setting accordingly very close to the actual temperatures near the membrane. Temperatures obtained from the measurements for the two higher power settings are biased by the self-absorption of the thermocouple, and need to be also corrected for the influence of strong local heat convection, which continuously supplies a stream of cold medium to the lipid assemblies on the surface. We estimate that the temperatures on the surface are likely not exceeding 70 and 90 °C for the two higher power settings. More accurate direct temperature determinations would be possible using ion conductivity measurements in a glass capillary[1], or a different means of temperature control (bulk heating) can be considered.

### S3. Image analyses

We performed image analysis using Matlab2018a to detect the compartments in the recorded micrographs to be able to count their number and to calculate their diameter, over time. Briefly, a suitable threshold to convert the gray scale micrographs to binary images, was adjusted every 250 frames of the time series to correct the intensity fluctuations, photo-bleaching and similar effects. Next, *imfindcircles* function was used to detect the circles using circular Hough transform. **Fig. S3** shows a sample output image from the analysis shown in **Fig. 3c**. A Matlab script has been provided as a separate supplementary file.

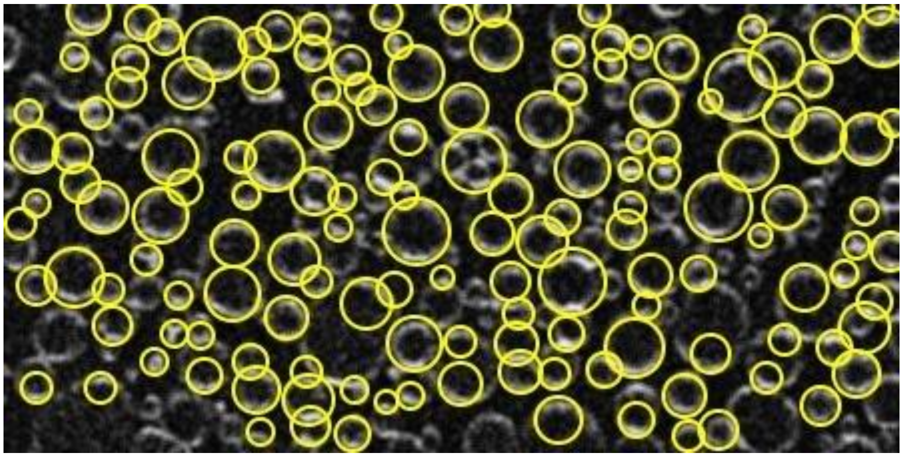

**Figure S3 |** Sample output image of *imfindcircles* function of Matlab, showing the identified circular compartments.

### S4. Details of computational model

We perform two sets of simulations: (1) fusion initiating at the equator of the compartments and (2) fusion initiating along the connecting tube. For both simulations the initial configurations are similar (schematically depicted in **Fig. S4**). In the first case, the vertices highlighted in red are connected along the contact (neck) line (red solid line in **Fig. S4**), and the circumference of the contact line is constrained. In the second case, two vertices with the same initial coordinates are placed along the red dashed line in **Fig. S4**, which allows us to define two faces that separate the two neighboring vesicles. In the set of simulations regarding case 1, the energy minimization leads to a separation of the vertices within the first steps, thus preventing an intersection of the two vesicles. In the second set of simulations the midline along the tube (red dashed line in **Fig. S4**) is constrained. In all simulations, the outer points (shown in green in **Fig. S4**) and the faces between them are constrained to a cylinder of radius 1, where we define all lengths in unites of the cylinder radius. The lowest vertices and the lines connecting these vertices are constrained to a height $z$=0, while all other vertices, lines and faces are constrained to $z$>0. The total area is constrained to 450.

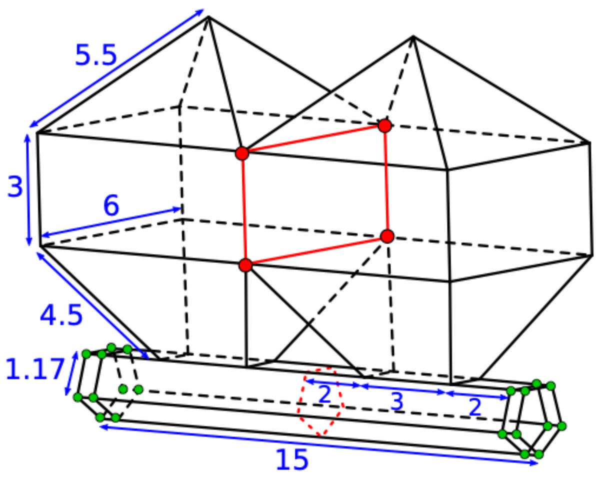


**Figure S4 | Initial configuration of the simulation.** All lengths are given in units of the tube radius.

We determined the bending energy by integrating the mean curvature over the entire surface, using the built-in *star_perp_sq_mean_curvature* method. The *Surface Evolver* software[2] uses finite element methods on a two-dimensional mesh in space to minimize the bending energy, which is defined as the integral of the squared mean curvature over the surface. The bending energy $E_v$ of each vertex reads:

$$E_v = A_v \frac{3}{4}\left(\frac{\nabla A_v \boldsymbol{N}_v}{\boldsymbol{N}_v \boldsymbol{N}_v}\right)^2, \quad (1)$$

with $A_v$ the area of the facets adjacent to the vertex and $\boldsymbol{N}_v$ the volume gradient, which is defined as:

$$\boldsymbol{N}_v = \frac{1}{6}(\boldsymbol{v}_1 \times \boldsymbol{v}_2 + \boldsymbol{v}_2 \times \boldsymbol{v}_3 + \cdots + \boldsymbol{v}_n \times \boldsymbol{v}_1), \quad (2)$$

with $\boldsymbol{v}_1, \ldots, \boldsymbol{v}_n$ the neighboring vertices. A detailed description of the numerical methods can be found in the *Surface Evolver* manual[3].

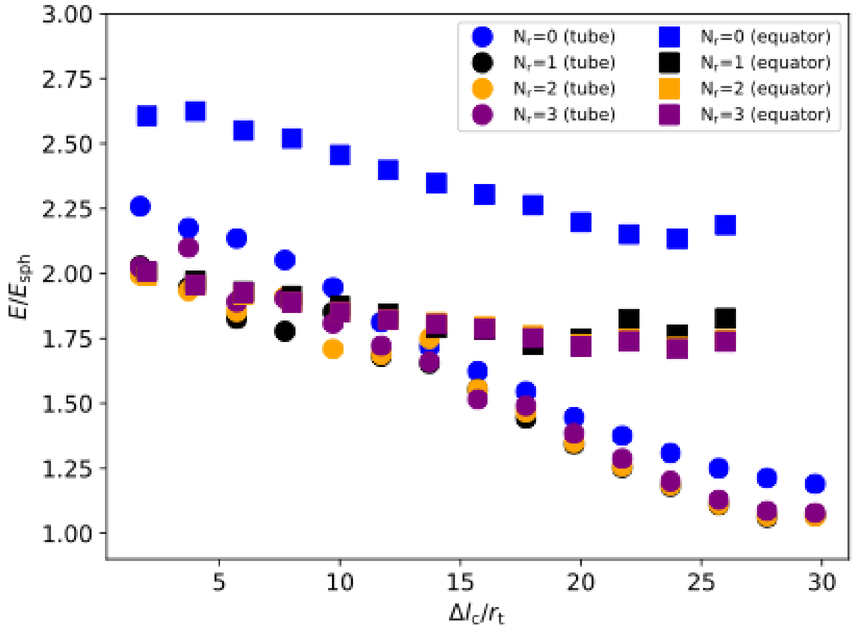


**Figure S5 | The bending energy, rescaled by the energy of a spherical vesicle is shown in dependence of the length of the contact line.** The number of mesh refinements $N_r$, corresponding to simulation step 1, 2, 3 and 4 are shown in different colors.

The equilibrated membrane shape is obtained through a series of four energy minimization steps, which read in the *Surface Evolver* letter code:

*Fusion at the tube:*

step 1: {g 20; u}3;{{V;g 20}30; u ;g 20}2
step 2: r;w 0.1;U;{V;g 50}10; U;g 20
step 3: r;w 0.05;U;{V;g 50}10; U;g 20
step 4: r;w 0.02;U;V;g 50;u;{V;g 50}9; U;g 20

*Fusion at the equator:*

step 1: g 50; u; g 100;U; g 100;U
step 2: r;w 0.1;{{u;g 50}4;{V;g 50}}2;g 20
step 3: r;w 0.05;{{u;g 50}4;{V;g 50}}2;g 20
step 4: r;w 0.02;{{u;g 50}4;{V;g 50}}2;g 20

Step 2, 3 and 4 start with a mesh refinement (‘r’). In **Fig. S5** we see that only the first mesh refinement leads to a significant improvement of the energy minimization. In **Fig. 4** in the main text the minimal energy found within the four simulation steps is shown.

### S5. Stable pore formation in fused compartments

The mathematical model for fusion shows the formation of a stable pore when fusion initiates at the equator of two adjacent compartments. We also observe stable pores in fused compartments in our experiments (**Fig. S6**, also **Fig. 3** of the main manuscript). Such pores are not observed before fusion, or if the fusion occurs between the compartments initially residing on the same nanotube (**Fig. 3d-k** for fused compartments initially were on the same nanotube vs. **Fig. 3l-u** for compartments initially were on different nanotubes).

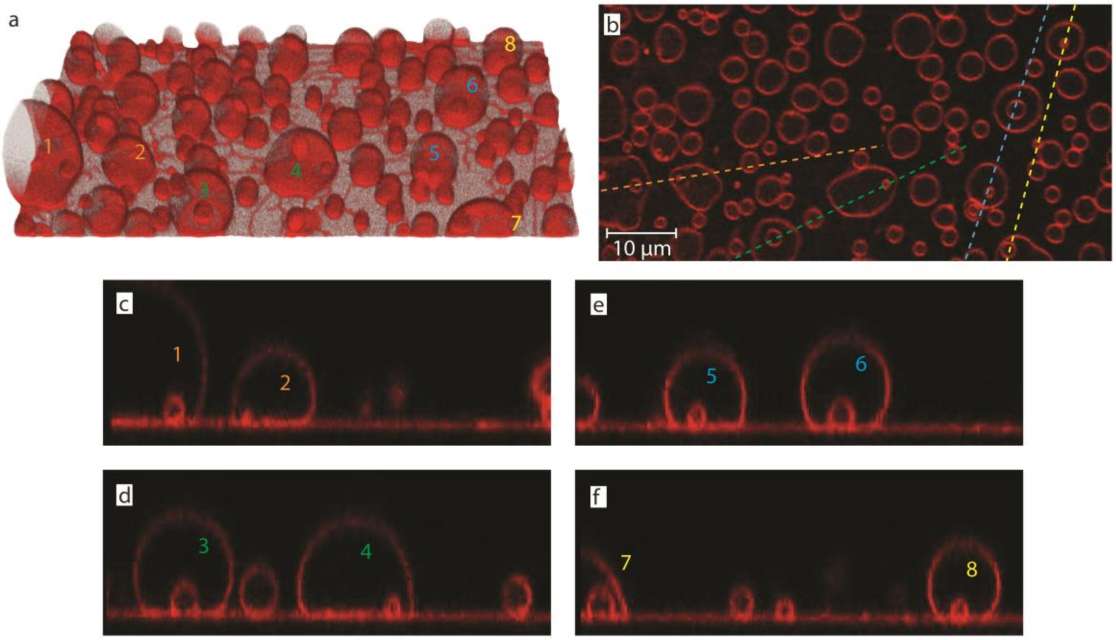


**Figure S6 | Stable pores in fused compartments. (a)** 3D confocal micrograph of the formed protocells **(b)** cross section of protocell sample close to the surface (x-y plane) **(c-f)** cross sections of protocell sample (x-z plane). Protocells numbered 1-8 in **(a)** have small cavities formed after fusion of adjacent protocells. Cross sectional profile of the numbered protocells, along the color-coded dashed lines in **(b)**, are shown in **(c-f)**.

### S6. Encapsulation of fluorescein vs. RNA inside the protocells

As described in the manuscript, we delivered the RNA to the compartments locally using an open-space microfluidic device which resulted in the encapsulation of RNA inside some of the compartments. FAM conjugated 10 base long polyA RNA oligonucleotides were prepared in nuclease-free water. For experiments, the RNA concentration was adjusted to 40 μM with HEPES buffer. The encapsulation efficiency of free fluorescein sodium salt is compared to FAM-RNA oligonucleotides. The fluorescein encapsulation shows higher efficiency (**Fig. S7**). Only a few protocells encapsulate and maintain RNA fragments. This might be due to the higher molecular weight and structure of RNA compared to the fluorescein dye.

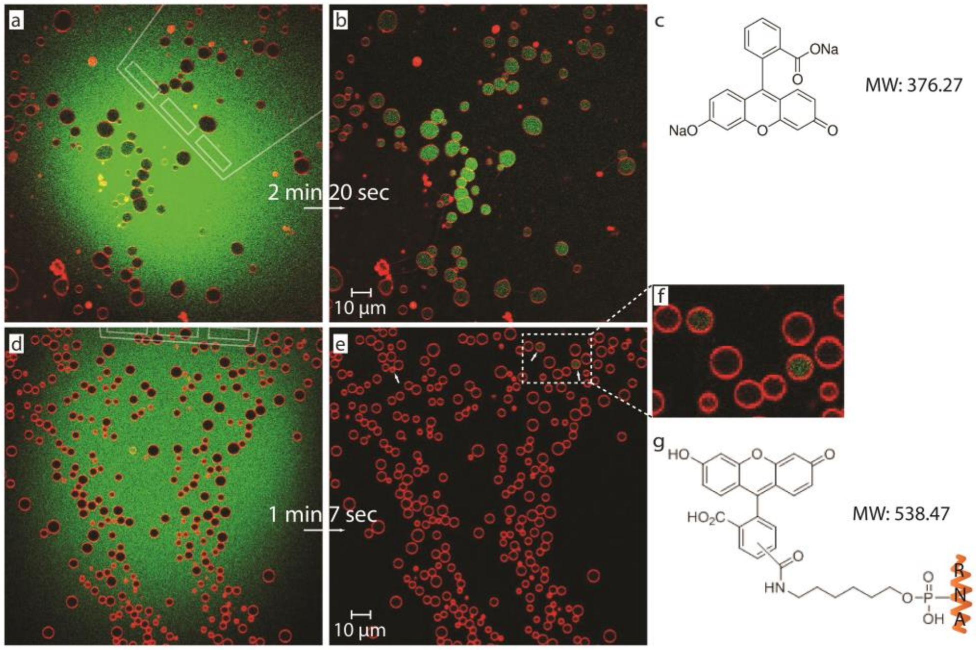


**Figure S7 | Encapsulation of fluorescein dye vs. FAM conjugated RNA oligonucleotides.** A microfluidic pipette is used for the superfusion of free fluorescein and FAM-conjugated RNA oligonucleotides to a membrane area populated with protocells. **(a)** Confocal micrograph of a membrane area with the microfluidic pipette re-circulating free fluorescein above it (top view). **(b)** After exposure is stopped, the initially fluorescein-free compartments are observed with fluorescein cargo. **(c)** fluorescein sodium salt molecule (MW: 376.27 g/mol). **(d)** Confocal micrograph of a membrane area with the microfluidic pipette re-circulating FAM-conjugated RNA fragments above it (top view). **(e)** After exposure, only a few protocells, shown in **(e)** (white arrows), and in **(f)**, encapsulate and maintain RNA fragments in the ambient aqueous solution. **(g)** Fluorophore-conjugated RNA molecule (MW: 538.47 g/mol).

### S7. Supplementary Movies

**Movie S1-3 | Rapid nucleation and growth of protocells.** Movies S1-S3 show the rapid nucleation and growth of vesicular compartments from the underlying lipid nanotube network upon increase in temperature.

The region shown in Movie S1 ‘Experiment 2’ corresponds to **Fig. 1c-e** and **Fig. 2a**. The region shown in Movie S1 ‘Experiment 1’ corresponds to **Fig. 1f-h**. ‘Experiment 1-2’ Movie S1 is accelerated 94x and 71x, respectively.

‘Experiment 3-5’ in Movie S2 are accelerated 101x, 114x and 136x, respectively.

The region shown in Movie S3 corresponds to **Fig. 2c-f**, and the movie is accelerated 5x.

**Movie S4-6 | Fusion of protocells.** Movie S4-S6 show the fusion of protocells on lipid nanotube networks upon exposure to higher intensity IR laser radiation.

Movie S4 and S5 correspond to **Fig. 3**. In Movie S5, the outline of lipid nanotube network is overlaid to facilitate the observation of location of fusion events with respect to the nanotubes, i.e. same or different tubes. Movies are sped up 5x. Between each part (Part I-III) the recording is stopped and re-started.

Movie S6 is accelerated 85x.

**Movie S7 | Encapsulation of RNA oligonucleotides and following fusion of protocells.** Movie S7 shows the superfusion of FAM-conjugated RNA oligonucleotides by means of a microfluidic pipette (Part I) followed by fusion events induced by activation of the IR laser (Part II).

Movie S7 Experiment 1 and Experiment 2 correspond to **Fig. 5c-m** and **Fig. 5r-t**, respectively. Movies are accelerated 5x.
